# A microfluidic platform for multi-marker profiling of extracellular vesicles from single-cell-derived clones

**DOI:** 10.64898/2026.03.13.711619

**Authors:** Junyoung Kim, Doru Petrisor, Dan Stoianovici, Sarah Amend, Kenneth Pienta, Chi-Ju Kim

## Abstract

Extracellular vesicles (EVs) carry molecular cargo that can reflect the real-time state of parental cells, yet most in vitro EV analyses rely on bulk approaches and therefore average over pronounced heterogeneity in both cell and EV populations. Here, we present a semi-open microfluidic platform that enables multi-marker profiling of EVs released from single-cell-derived clones, allowing EV signatures to be linked to clonal progeny originating from a single parental cell. The platform integrates aligned cell and EV arrays containing 17,305 wells, assembled with a 3D-printed housing to capture released EVs in one-to-one matched wells. Captured EVs are immunolabeled for canonical tetraspanin markers (CD9, CD63, CD81) and EpCAM, imaged by high-resolution fluorescence microscopy, and quantified using an automated image-analysis pipeline. Applying the platform to single-cell-derived PC3 clones revealed substantial heterogeneity in EV marker co-expression, with hierarchical clustering identifying four distinct tetraspanin co-expression profiles. The fraction of EpCAM-positive EVs increased with PC3 cell proliferation, as assessed by endpoint cell number, whereas free (non-EV-associated) EpCAM showed no correlation. This platform enables near single-EV-level, multi-marker profiling from single-cell lineages and provides a practical approach to simultaneously dissect both cellular and EV heterogeneity.

## Introduction

Extracellular vesicles (EVs) are nanoscale particles released by cells that mediate intercellular communication.^1–5^ EVs are released through two major biogenesis routes: (1) exosomes are released when MVBs/MVEs fuse with the plasma membrane, and (2) microvesicles/ectosomes are shed via outward budding from the plasma membrane.^1,2,4,5^ Consequently, EV composition and their secretion can vary with cellular state and microenvironmental cues,^1,4^ thereby shaping intercellular communication through the transfer of proteins, lipids, and nucleic acids.^1,2,4,5^

Accordingly, EVs can naturally carry diverse molecular signatures from their parental cells, reflecting real-time cellular status.^4,5^ Interestingly, our previous finding showed that EV cargo not only mirrors the parental cellular state,^6^ but may also paradoxically include unwanted or waste molecules generated during cellular activity that may be exported to maintain parental cell phenotypes.^1,6–9^ Therefore, investigating EV release pathways and the cargo that EVs carry is worthwhile for precisely revealing the molecular signatures of cellular status.

EVs are typically obtained from cell culture by collecting cell culture conditioned medium (CCM) in most in vitro studies.^10^ This approach yields EV samples that represent pooled vesicles released from millions of cells. However, substantial cellular heterogeneity exists even within the same cell type, and EV subpopulations are also heterogeneous.^4,10,11^ If EVs released from individual cells can be analyzed separately, it becomes possible to decipher EV molecular signatures in direct association with their parental cells, enabling discoveries that are not accessible through conventional bulk approaches.

Several robust microfluidic devices have been developed to study EVs secreted from single cells,^12–17^ enabling single-EV level detection,^16,17^ quantification of EV secretion rates,^17^ and analysis of tumor patient samples.^12,13^ However, many of these platforms are not sufficiently flexible to capture diverse EV subpopulations, may not reliably support single-EV level detection, and/or require external equipment for device operation. Here, we developed a semi-open microfluidic system that enables single-cell trapping, EV capture regardless of EV subtypes, near single-EV level detection, and multi-marker profiling of EVs (**Figure 1**).

**Figure 1.**
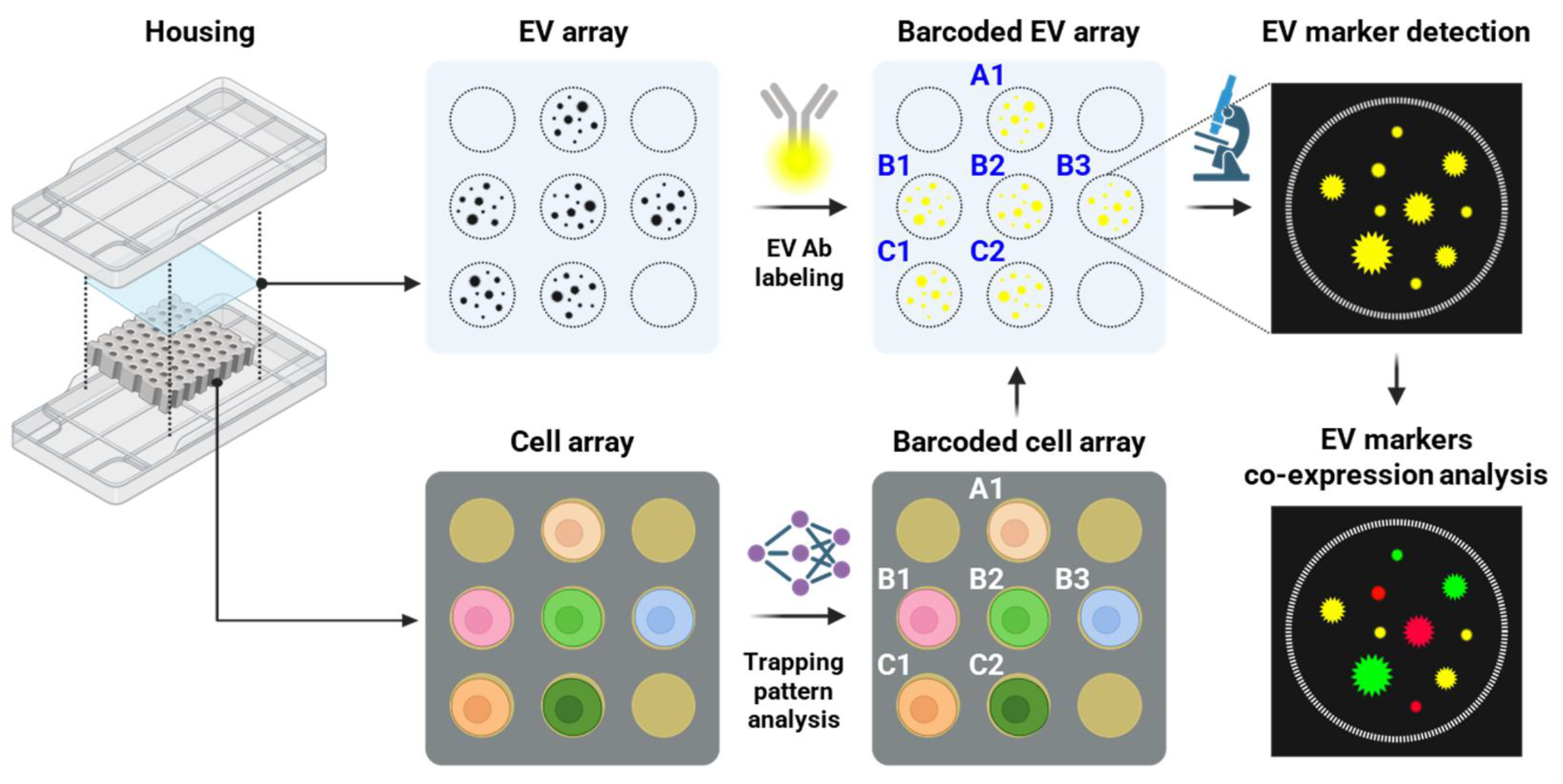
Schematic illustration of a single-cell-derived clonal EV analysis system. The system consists of an EV array, a cell array, and a housing module for aligned, secure assembly. Single cells are trapped in the cell array, and the resulting trapping pattern serves as a spatial “barcode” that enables one-to-one mapping of each cell-containing well to the corresponding well in the EV array. EVs released from single-cell-derived clones are captured on the EV array assembled above the cell array within the housing. EV surface markers and cargo expression are detected by immunofluorescence (IF) imaging using high-resolution microscopy. Expression of multiple EV markers and EV cargo is further evaluated by post-imaging co-expression analysis.

## Methods

### Validation of the EV-array detection method

For EV-array validation, PC3 cells were maintained in RPMI 1640 medium (Thermo Fisher Scientific, USA) containing 5 U/mL penicillin–streptomycin (PenStrep; Thermo Fisher Scientific, USA) and 10% exosome-depleted FBS (VWR, USA). PC3 cells were seeded at approximately 30% confluency and maintained for 48 h in a humidified tissue culture incubator (37 °C, 5% CO_2_). CCM was then collected and immediately pre-processed as previously described.^10^

To immobilize EVs on a substrate, 35 mm glass-bottom dishes (ibidi, USA) were coated overnight at 4 °C with PDL (0.1 mg/mL; Gibco, USA) and rinsed three times with D.I. water. Pre-processed CCM was then applied to the PDL-coated surfaces and incubated overnight at 4 °C. Following three PBS washes, non-specific binding was blocked using ELISA Ultrablock (Bio-Rad, USA) for 1 h at RT.

For EV marker co-expression analysis, primary antibodies were diluted in blocking buffer and applied to the EV-immobilized surfaces at the following concentrations: PE mouse anti-human CD9 (1:100, BD Biosciences, Cat No. 341647, USA), Alexa Fluor® 488 anti-human CD63 (1:400, BioLegend, Cat No. 353038, USA), and APC mouse anti-human CD81 (1:100, BD Biosciences, Cat No. 551112, USA). Samples were incubated with antibodies at 4 °C overnight, followed by three washes with 1X PBS.

Fluorescently labeled EVs on the EV array were imaged using a Nikon Eclipse Ti2 microscope equipped with a DS-Qi2 camera and NIS-Elements software (v5.11) using tile acquisition and a Plan Apo λ 100X oil-immersion objective (Nikon Inc.). Excitation/emission settings were 480/535 nm for Alexa Fluor® 488 and FITC (green), 570/645.5 nm for PE (red), and 620/700 nm for APC (far-red). Images were analyzed using CellProfiler 4 software^18^ to quantify EV counts and evaluate EV markers co-localization.

### Device fabrication

### Cell-array

A PDMS-based cell trapping array (cell-array) containing circular trapping wells (100 µm diameter, 190 µm depth; 17,305 wells total) was fabricated by soft lithography using a master mold generated by photolithography at the Johns Hopkins University (JHU) WSE Whitaker Microfabrication Lab (**Supplementary Figure S1**). Fabricated PDMS devices were washed and sterilized with 100% isopropanol (IPA), then fully dried in a biosafety cabinet (BSC) under UV exposure for 30 min. The cell-array surface was treated using a corona plasma treater (BD-20AC Laboratory Corona Treater, Electro-Technic Products, USA) for 1 min 30 s, coated with 0.1 mg/mL PDL at 37 °C overnight, and washed three times with 1X PBS.

### EV-array

PDMS base and curing agent were mixed at a 10:1 ratio (base:curing agent), poured into a 150 mm dish, and cured at 65 °C for 4 h. PDMS square blocks (approximately 22 mm X 22 mm) were cut from the cured PDMS slab. The EV-array blocks were washed and sterilized with 100% ethanol (EtOH) and dried in a BSC under UV exposure for 30 min. Blocks were treated with a corona plasma treater for 1 min 30 s, coated with 0.1 mg/mL PDL at 37 °C overnight, and washed three times with D.I. water.

### Housing

Housings used to assemble the cell-array and EV-array were fabricated by 3D printing at the JHU Urology Robotics Lab (**Supplementary Figure S2**). The housing consists of top and bottom frames incorporating permanent magnets, enabling alignment and secure assembly of the cell-array and EV-array. This magnetic clamping, together with the elastic deformability of PDMS, allows leak-free assembly. Two housing designs were used: (i) a compact housing (**Supplementary Figure S2a**) for routine culture and (ii) a specialized housing (**Supplementary Figure S2b**) designed to facilitate visualization of cellular behaviors during culture in the assembled device. Printed housings were washed and sterilized with 100% EtOH and dried in a BSC under UV exposure for 30 min.

### Device operation

### Cell trapping and pre-culture on the cell-array

PC3 cells were cultured in RPMI 1640 supplemented with 10% FBS and 5 U/mL PenStrep in a humidified tissue culture incubator (37 °C, 5% CO_2_). Cells were resuspended in 500 µL of culture medium and dispensed onto the PDL-coated cell-array to uniformly cover the well region, followed by a 5 min settling period. Untrapped cells located between wells were removed by gently squeegeeing the surface twice with a glass coverslip in two orthogonal directions. The cell-array containing trapped cells was transferred to a 60 mm dish and supplied with 5 mL of medium.

The entire cell-array area was imaged using an EVOS M7000 microscope with tile acquisition to identify wells initially containing a single trapped cell. Arrays were then cultured for 48 h or 72 h in a humidified tissue culture incubator (37 °C, 5% CO_2_). After culture, the entire cell-array was re-imaged using EVOS M7000 tile acquisition to identify “active” wells containing viable cells immediately prior to housing-based device assembly.

### Device assembly and culture in the housing

The assembled device consists of a housing-bottom frame with magnets, a glass slide (75 mm X 25 mm), the cell-array, the EV-array, and a housing-top frame with magnets. Briefly, a sterilized glass slide was placed on the housing-bottom frame. To minimize background signal from residual EVs from medium components, the cell-array was washed three times with RPMI 1640 supplemented with 10% exosome-depleted FBS and 5 U/mL PenStrep. The washed cell-array was placed on the glass slide, and the EV-array was then positioned to cover the cell-array. The housing-top frame was assembled onto the housing-bottom frame via magnetic clamping. The assembled device was placed in a 150 mm dish, and sterile water was added to the dish to mitigate evaporation of culture medium within individual trapping wells. Devices were incubated for 24 h in a humidified tissue culture incubator (37 °C, 5% CO_2_).

After 24 h, the device was disassembled. The EV-array was immediately transferred to a dish containing 1X PBS to prevent drying. The cell-array was immediately stained with Hoechst 33342 Ready Flow™ Reagent (Invitrogen, USA) for 30 min in a humidified tissue culture incubator (37 °C, 5% CO_2_), washed three times with medium, and imaged using EVOS M7000 tile acquisition to determine active wells containing viable (analyzable) cells.

### Immunolabeling and imaging of EVs on the EV-array

EV-arrays were washed three times with 1X PBS and blocked with blocking buffer for 1 h at RT. Primary antibodies were diluted in blocking buffer and applied as follows. For EV marker co-expression analysis: PE mouse anti-human CD9 (BD Biosciences, 341647, 1:100), Alexa Fluor® 488 anti-human CD63 (clone H5C6; BioLegend, 353038, 1:400), and APC mouse anti-human CD81 (BD Biosciences, 551112, 1:100). For EV-EpCAM expression analysis: FITC mouse anti-human CD9 (BD Biosciences, 341646, 1:100), Alexa Fluor® 488 anti-human CD63 (clone H5C6; BioLegend, 353038, 1:400), FITC mouse anti-human CD81 (BD Biosciences, 551108, 1:100), and PE anti-human CD326 (EpCAM) (BioLegend, 324206, 1:400).

Samples were incubated with antibody mixtures at 4 °C overnight, followed by three washes with 1X PBS. Fluorescently labeled EVs were imaged on a Nikon Eclipse Ti2 microscope using tile acquisition and a Plan Apo λ 100X oil objective. Excitation/emission settings were 480/535 nm for Alexa Fluor® 488 and FITC (green), 570/645.5 nm for PE (red), and 620/700 nm for APC (far-red). Images were analyzed using CellProfiler 4 software^18^ to quantify EV counts and assess co-localization among EV markers and EpCAM.

### Image-based cell viability assay

To assess cell viability after culture in the assembled device, immediately after device disassembly, trapped cells in the cell-array were stained with 2 µM calcein AM (Invitrogen, USA) and Hoechst 33342 Ready Flow™ Reagent for 30 min in a humidified tissue culture incubator (37 °C, 5% CO_2_). The array was washed three times with medium and imaged directly using an EVOS M7000 microscope with tile acquisition.

### EV markers (CD9, CD63, and CD81) co-expression analysis

For EV marker co-expression analysis, post-trapping culture on the cell-array was conducted for 48 h. A total of 33 candidate “positive” wells satisfying both criteria were selected: (i) wells containing a single trapped cell immediately after trapping and (ii) wells containing a viable cell after culture in the assembled device with the EV-array. In addition, 16 negative-control wells containing no cells were used to define background thresholds. For each marker (CD9, CD63, and CD81), the positivity threshold was defined as the mean + 3 standard deviations (SD) of the corresponding marker counts measured in the 16 negative-control wells. A well was classified as positive only if all three marker counts exceeded their respective thresholds (AND rule).

Among the analyzed wells, the EV marker co-expression percentages from the 17 wells that passed the positivity criteria were transformed into z-scores. These z-scores were used to generate heatmaps and perform hierarchical clustering in Morpheus (Broad Institute) using the following settings; metric: one minus Pearson correlation; linkage method: average; clustering axis: columns (single cells).

### EpCAM and EV marker co-expression analysis

For EpCAM and EV marker co-expression analysis, post-trapping culture on the cell-array was conducted for 72 h. Using the same selection criteria as those used for the EV marker co-expression analysis, 37 candidate positive wells were identified (single-cell occupancy after trapping and viability after culture in the assembled device). In addition, 20 negative-control wells containing no cells were used to define background thresholds. For EV counts and EpCAM counts, positivity thresholds were defined as the mean + 3 SD of the corresponding counts measured in the 20 negative-control wells. A well was classified as positive only if the relevant marker counts exceeded their respective thresholds (AND rule).

For the 35 wells that passed the positivity criteria, (i) the percentages of EV marker and EpCAM co-expression out of total EVs and (ii) the percentage of free EpCAM out of total EpCAM were transformed into z-scores and visualized as heatmaps using Morpheus.

## Results

### Development and validation of the EV array

We developed a method for detecting EV markers (CD9, CD63, CD81) and analyzing their co-expression by combining EV immunofluorescence (EV-IF) imaging with automated EV quantification during post-image processing (**Figure 2a**). To validate this EV detection strategy, we first evaluated it using a bulk EV preparation. PC3 cell–derived EVs were immobilized on a PDL-coated 35-mm glass-bottom dish. Fluorophore-conjugated antibodies were then applied to label tetraspanin EV markers (CD9, CD63, CD81).

**Figure 2.**
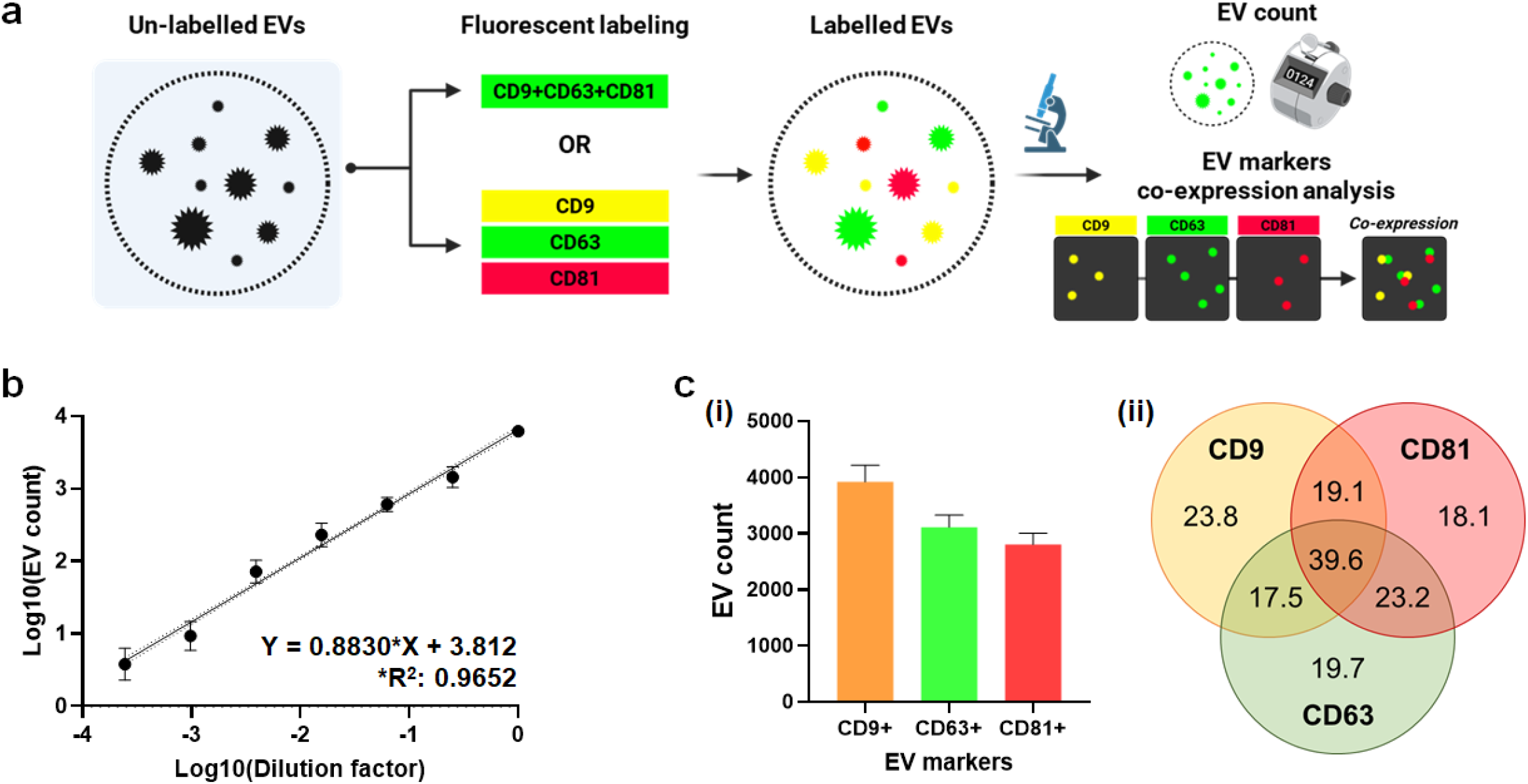
Validation of the EV array. **a. Overall workflow of EV array labeling and analysis**. The EV array was labelled either with an antibody cocktail of EV markers (CD9, CD63, CD81) using a single color fluorophore (green), or with the markers conjugated to three spectrally distinct fluorophores. EVs positive for CD9, CD63, and/or CD81 and their marker co-expression were quantified using CellProfiler™ 4, **b. Quantification of EV counts as a function of PC3 CCM dilution**. EVs were labelled with an antibody cocktail (CD9, CD63, CD81) conjugated to the single color fluorophore (green). EVs were counted within a virtual, centered 100-µm-diameter ROI to match the effective well area analyzed in the fully integrated system. Images containing severe artifacts (e.g., large dust particles) that obscure true EV signals (particles below approximately 1 µm) were excluded prior to quantification. The numbers of images analyzed by CellProfiler were: no dilution (×1.00E0), 30; ×2.50E−1, 37; ×6.25E−2, 35; ×1.56E−2, 37; ×3.91E−3, 32; ×9.77E−4, 30; ×2.44E−4, 31; ×6.10E−5, 30, **c. Co-expression analysis of EV markers (CD9, CD63, CD81)**. EVs were labelled with three fluorophore-conjugated antibodies (CD9, yellow; CD63, green; CD81, red). EVs were quantified within the same virtual, centered 100-µm-diameter ROI for consistency. A total of 35 images were analyzed by CellProfiler. Panel (i) shows the number of EVs positive for each marker, and panel (ii) shows a Venn diagram summarizing marker co-expression as percentages.

To quantify EV counts as a function of the PC3 CCM dilution factor, we used an antibody cocktail of green fluorophore–conjugated EV marker antibodies (FITC: CD9 and CD81; Alexa 488: CD63), and fluorescence signals were acquired using high-resolution microscopy. EV counts decreased proportionally with increasing dilution of PC3 CCM across seven concentrations including undiluted CCM. Quantitative analysis showed a wide dynamic range and strong linearity between EV counts and dilution factor (R^2^ = 0.9652; **Figure 2b**).

To validate the EV marker co-expression analysis, we used three spectrally distinct fluorophore-conjugated antibodies (CD9-PE, CD63–Alexa 488, and CD81-APC). Signals from the three fluorophores were acquired by high-resolution microscopy and quantified during post-image processing using a CellProfiler 4 pipeline. The results indicate that the mean (±SD) counts of EVs positive for CD9, CD63, and CD81 were 3918 (±300), 3109 (±216), and 2801 (±200), respectively (**Figure 2c-i**). EV marker co-expression percentages and counts are summarized in **Figure 2c-ii** and **Figure S3**, showing that the percentages of CD9+CD63+ EVs, CD9+CD81+ EVs, CD63+CD81+ EVs, and CD9+CD63+CD81+ EVs were 57.1% (±1.8%), 58.7% (±1.4%), 62.8% (±2.0%), and 39.6% (±1.9%), respectively. Together, these validation assays demonstrate robust quantitative EV counting and reliable co-expression profiling, supporting application of this EV marker detection strategy to single-cell-associated EV marker profiling.

### Development and validation of the cell array

We developed the cell array workflow by integrating microwell-based cell trapping and verifying each step using whole-array imaging (**Figure 3a**). Briefly, PC3 cells were trapped in fabricated microwells (100 µm diameter) using a seeding-and-squeegeeing method. The cell array was then covered with the EV array and assembled using a 3D-printed housing. Trapped PC3 cells were cultured for 48 h to expand from a single parental cell and to ensure sufficient EV release for downstream analysis.

**Figure 3.**
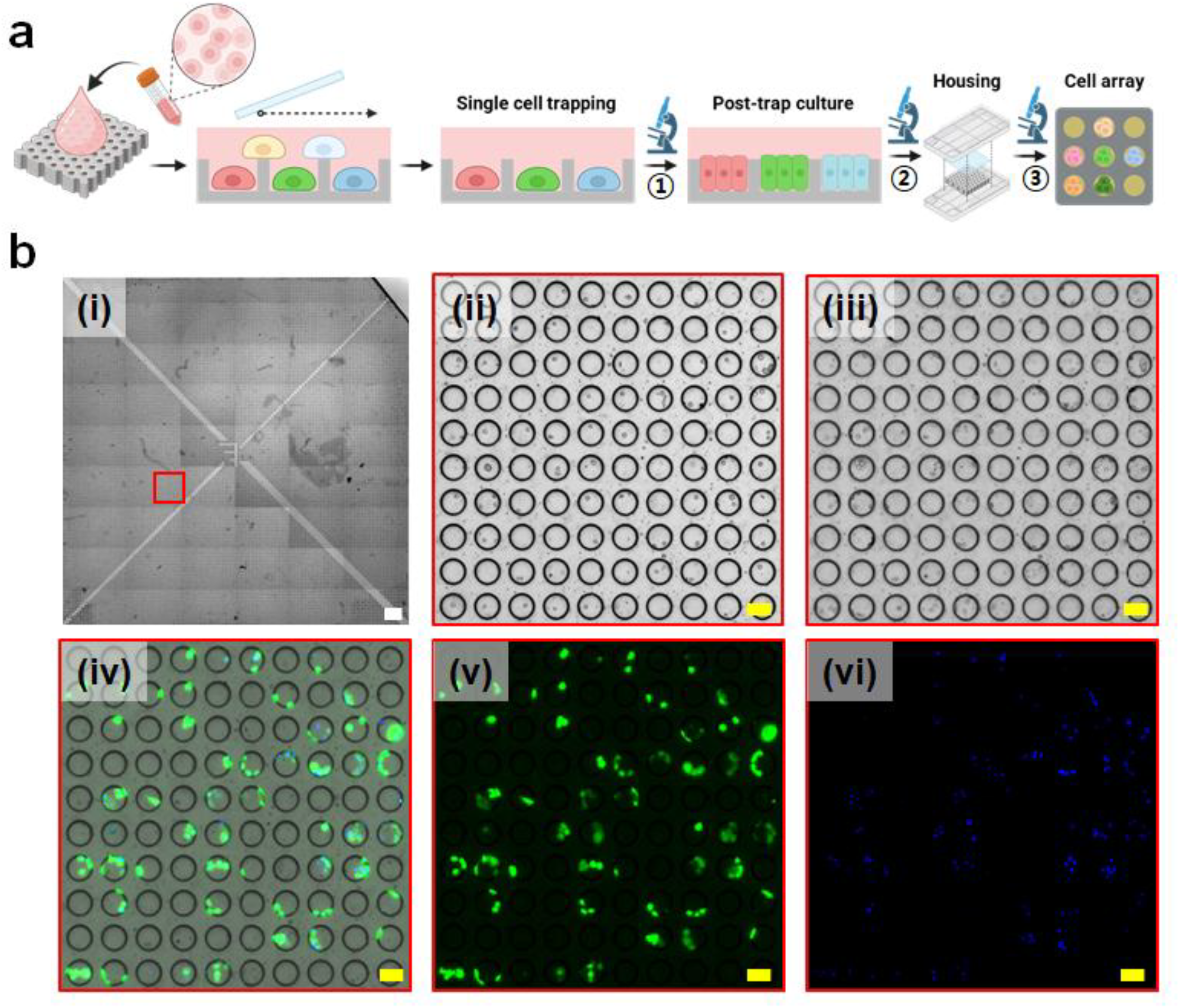
Validation of the cell array. **a. Schematic workflow of the cell array operation**. PC3 cells in suspension were uniformly loaded onto the cell array wells. Excess cells were removed by gently squeegeeing the array surface with a coverslip, followed by imaging (1). The cell array was then incubated for 48 h after trapping and imaged again (2). Next, the cell array and EV array were assembled within the housing and cultured for an additional 24 h, followed by imaging (3). **b. Representative images of cell trapping and viability assay after 24 h of on-trap culture within the housing**. (i) Whole-field image of the cell array before housing. Magnified views (red box) show (ii) the cell array immediately after trapping, (iii) the cell array immediately before housing assembly, and (iv–vi) the cell viability assay after culture in the assembled device: (iv) merged overlay, (v) Calcein-AM, and (vi) Hoechst. Scale bars: whole-field view (white), 1,000 µm; magnified view (yellow), 100 µm.

The cell array was imaged immediately after trapping (**Figure 3b-ii**), before housing (**Figure 3b-iii**), and after culture (**Figure 3b-iv**), demonstrating that cells remained viable and proliferated within the microwells. In addition, cells were labeled with Calcein AM to confirm viability after housing and culture (**Figure 3c-iv-vi**), showing that most cells were viable. These results indicate that the trapping-housing-culture workflow is suitable for maintaining cells and analyzing EVs released from trapped cells. Suitability of the workflow was further confirmed by time-lapse imaging using a customized open-window housing (**Supplementary Figure S2**) with CellTracker Green labeling (**Supplementary Video 1**), which confirmed active cell proliferation (**Supplementary Figure S4**).

### Matching the EV array with the cell array

To analyze EVs released from single-cell-derived clones, each well in the EV array was matched to the corresponding well in the cell array (**Figure 4a**). Cell array images acquired immediately after trapping (**Figure 4a-(1)**), before housing (**Figure 4a-(2)**), and after culture (**Figure 4a-(3)**) were used to identify analyzable wells, defined as wells that contained a single trapped cell at trapping and remained viable after culture. Wells lacking a trapped cell or showing loss of viability were excluded from downstream EV analysis.

**Figure 4.**
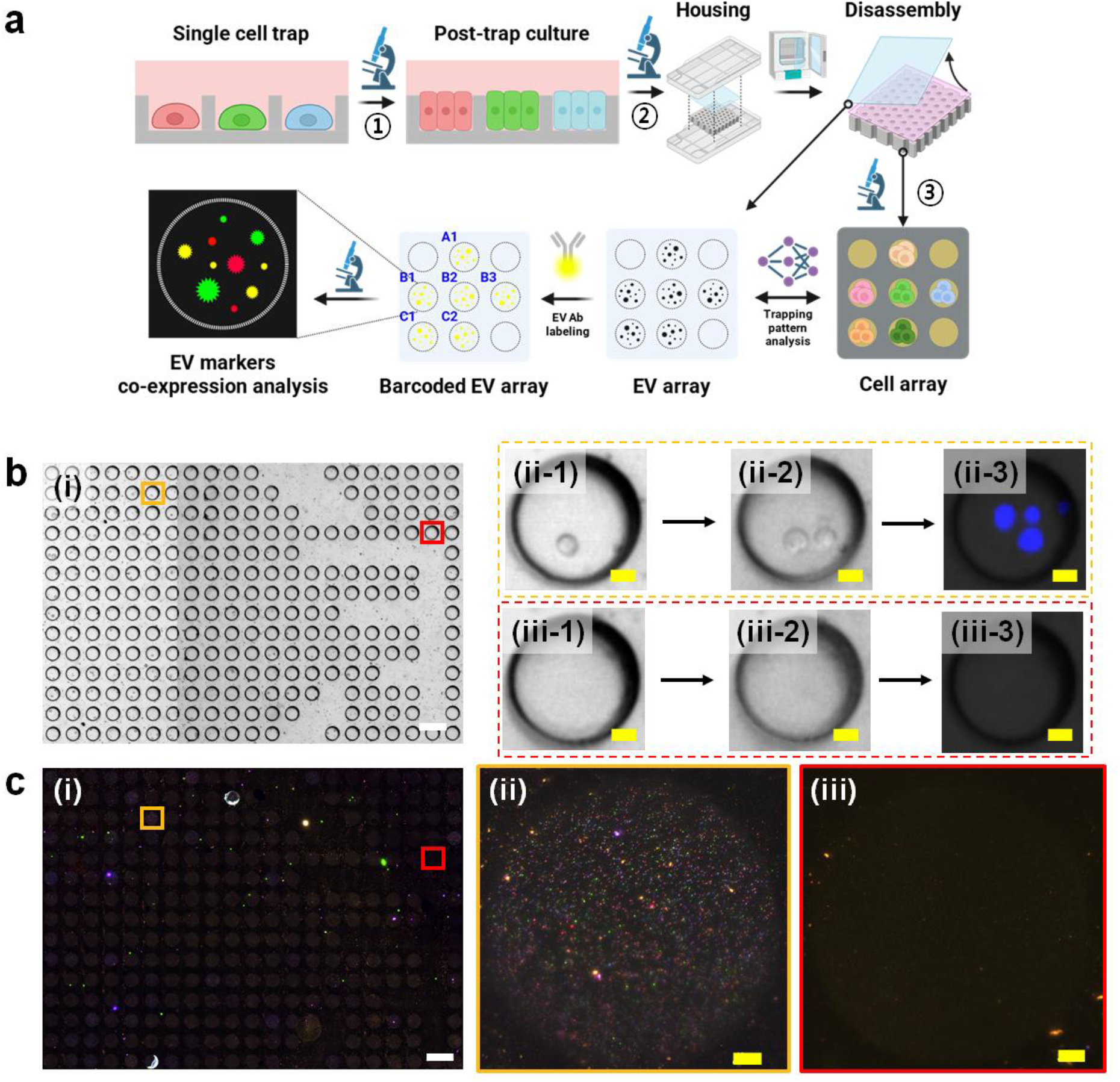
Matching between the cell array and the EV array. **a. Schematic workflow for matching the cell array and the EV array**. Single PC3 cells were trapped in the cell array and imaged (1). The cell array was then incubated for 48 h after trapping and imaged again (2). Next, the cell array and EV array were assembled within the housing for culture. After 24 h of culture, the arrays were disassembled and the cell array was imaged (3). The single-cell trapping pattern served as a spatial barcode to enable one-to-one mapping of each cell array well to the corresponding EV array well for subsequent EV marker expression analysis. EVs captured on the EV array were labelled for EV markers (CD9, CD63, CD81), and EV markers co-expression was quantified using CellProfiler™ 4. **b. Representative images of the cell array**. (i) Representative region of the cell array. (ii) A well (yellow box) initially trapping a single cell: (ii-1) after trapping, (ii-2) before housing assembly, and (ii-3) after culture. (iii) An empty well (red box): (iii-1) after seeding, (iii-2) before housing assembly, and (iii-3) after culture. Scale bars: (i) 200 µm; (ii, iii) 20 µm. **c. Images of the matched EV array**. (i) Tiled immunofluorescence (IF) image of the EV array (magnified view). Magnified views show EV marker signals from (ii) the EV array well corresponding to the single-cell-containing well (yellow box in panel b) and (iii) the EV array well corresponding to the empty well (red box in panel b). Scale bars: (i) 200 µm; (ii, iii) 10 µm.

Based on the image-based identification of analyzable wells of the cell array, the EV array image was spatially registered to the corresponding cell array image to establish one-to-one well correspondence. **Figure 4b-i** shows a representative region of the cell array after trapping, which was used to identify wells containing a single cell to ensure derivation from a single parental cell. Selected positive wells were further confirmed for cell viability primarily using the post-culture image and, secondarily, the pre-housing image. **Figure 4b-ii (yellow box)** indicates a selected positive well, whereas **Figure 4b-iii (red box)** shows a negative well without a trapped cell. **Figure 4c-i** shows the corresponding region of the EV array aligned to the cell array region in **Figure 4b-i**. A magnified view of the EV well for the selected positive well (**Figure 4c-ii**) visualizes the expression of three EV markers (CD9: blue, CD63: green, CD81: red) and demonstrates capture of EVs released from the trapped cells. In contrast, no appreciable EV marker signal was detected in the negative well (**Figure 4c-iii**). This matching strategy enables reliable analysis of EV markers and EpCAM expression in EVs released from cells originating from the same single parental cell.

### EV marker co-expression analysis

Cells derived from single parental cells could be clustered into four groups based on EV marker (CD9, CD63, CD81) co-expression profiles (**Figure 5, Supplementary Figure S5**). Features used for hierarchical clustering included the percentages of single-positive EVs (CD9+CD63-CD81-, CD9-CD63+CD81-, CD9-CD63-CD81+), double-positive EVs (CD9+CD63+CD81-, CD9+CD63-CD81+, CD9-CD63+CD81+), and triple-positive EVs (CD9+CD63+CD81+) (**Table 1**). Distinct co-expression patterns were also evident in Venn diagrams (**Figure S6**). These patterns were independent of the final cell number at the endpoint, which may indicate that proliferative phenotype may not influence the release of EVs expressing tetraspanin markers.

**Table 1.**
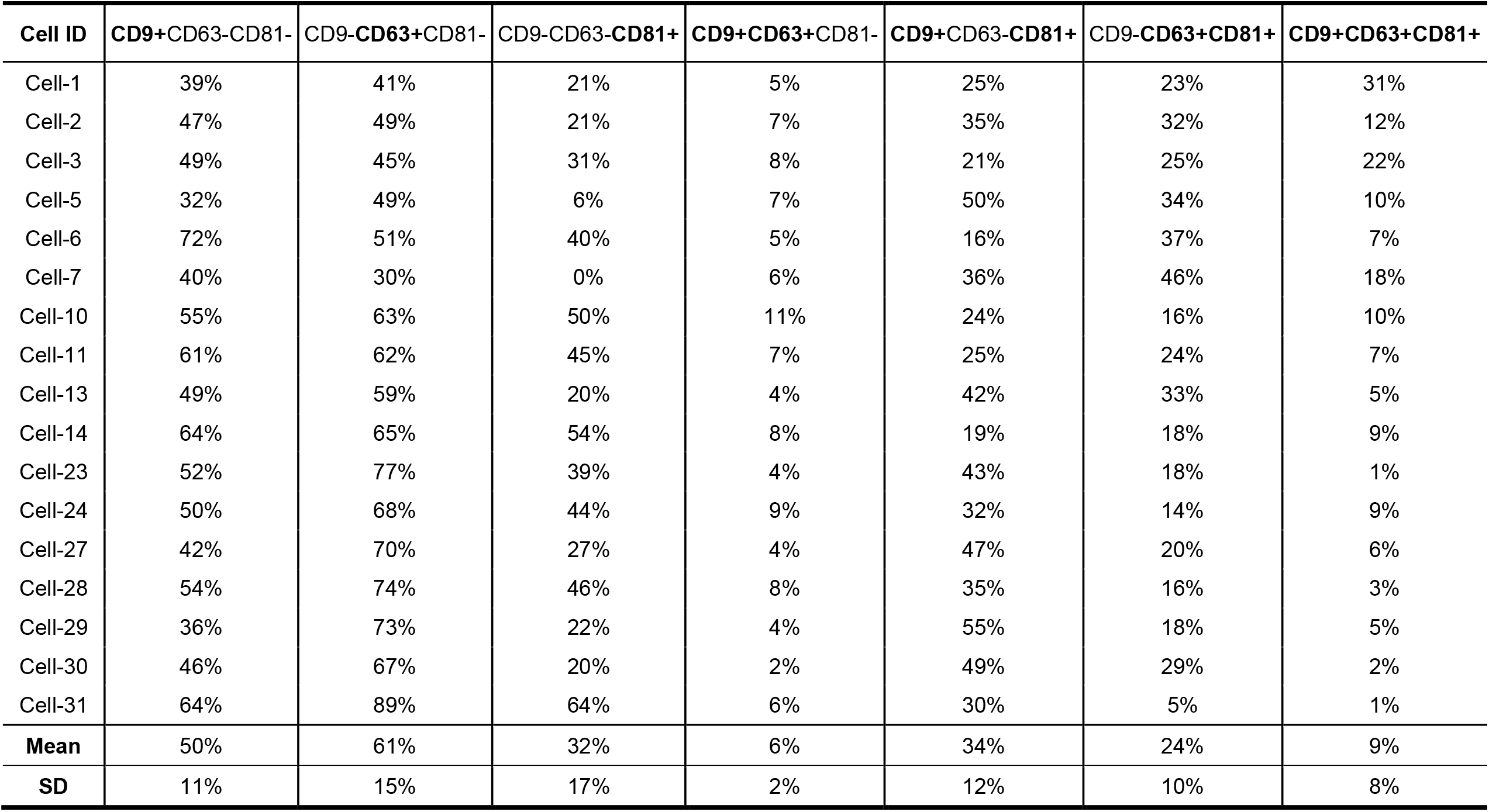
Percentages of EV-marker co-expression among EVs released from PC3 single-cell-derived clones.

**Figure 5.**
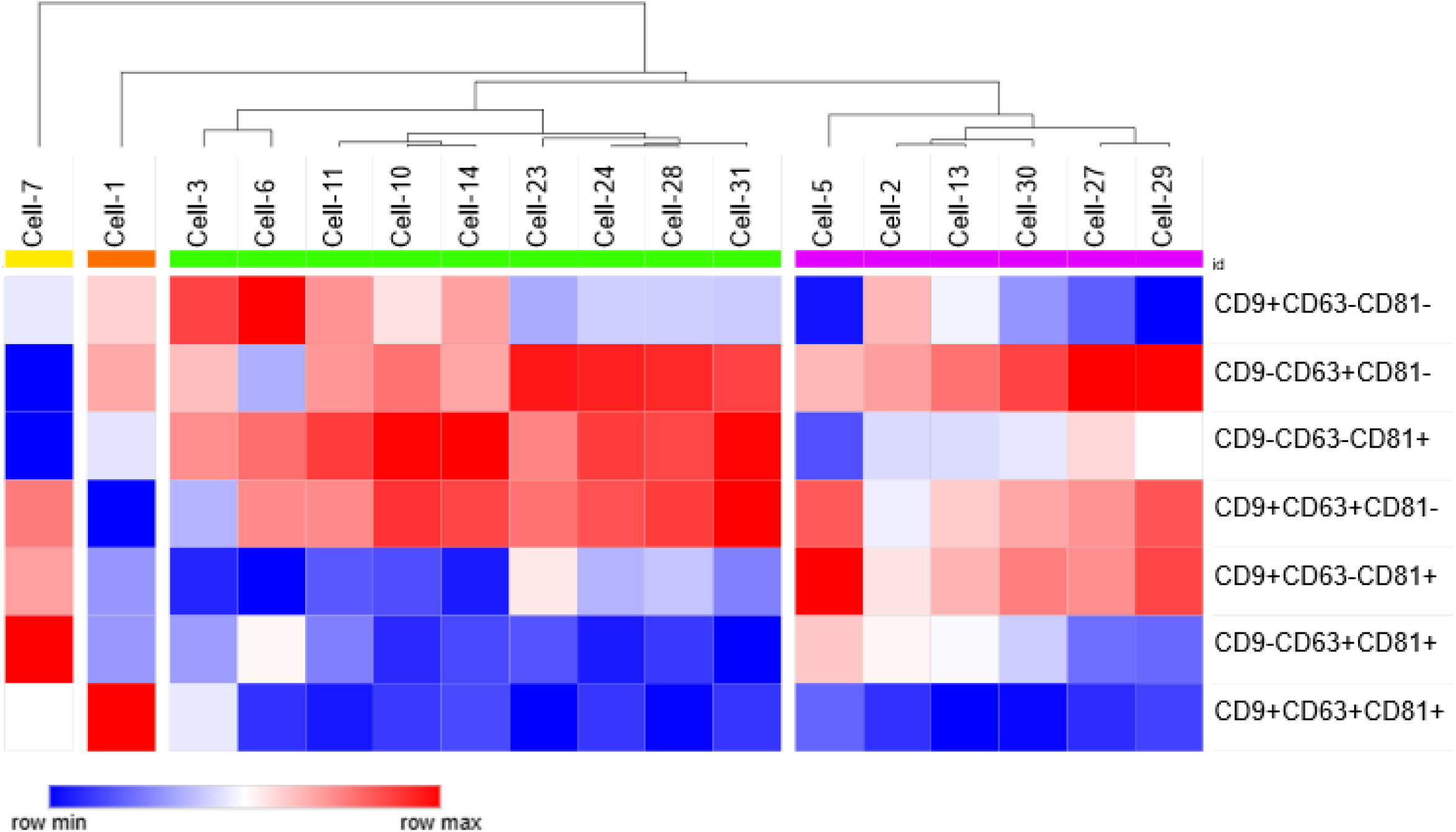
Heatmap of EV marker co-expression and hierarchical clustering. EV marker expression (CD9, CD63, CD81) in positive wells after thresholding was detected by high-resolution fluorescence microscopy, and EV marker co-expression levels were quantified using CellProfiler™ 4. Co-expression percentages were transformed into z-scores, which were used for heatmap visualization and hierarchical clustering in Morpheus (Broad Institute, https://software.broadinstitute.org/morpheus). Clustering was performed on columns (single cells) using one minus Pearson correlation as the distance metric and average linkage.

### Quantification of EpCAM co-expressed EVs

Using the developed system, we quantified EpCAM co-expressed EVs (EpCAM+ EVs) and free EpCAM released from single-cell-derived clones (**Figure 6, Table 2, Supplementary Figure S7)**. A heatmap comparing the percentage of EpCAM(+) EVs and the percentage of free EpCAM showed no correlation between the two (**Figure 6a**), suggesting that EpCAM loading into EVs may not be directly coupled to the level of remaining EpCAM. Interestingly, the percentage of EpCAM(+) EVs was positively correlated with the final cell number at the endpoint, whereas the percentage of free EpCAM showed no correlation (**Figure 6b**).

**Table 2.** Percentages of EV marker-EpCAM co-expression (among total EVs) and free EpCAM (among total EpCAM signal) in EVs released from PC3 single-cell-derived clones.

| Cell ID | EpCAM(+) EVs per Total EVs | Free EpCAM per Total EpCAM |
| --- | --- | --- |
| Cell-1 | 45% | 77% |
| Cell-2 | 28% | 73% |
| Cell-3 | 25% | 51% |
| Cell-4 | 46% | 69% |
| Cell-5 | 51% | 86% |
| Cell-6 | 55% | 81% |
| Cell-7 | 45% | 61% |
| Cell-9 | 41% | 76% |
| Cell-10 | 37% | 81% |
| Cell-11 | 35% | 79% |
| Cell-12 | 49% | 69% |
| Cell-13 | 48% | 71% |
| Cell-14 | 33% | 83% |
| Cell-15 | 25% | 77% |
| Cell-16 | 43% | 72% |
| Cell-17 | 39% | 59% |
| Cell-18 | 42% | 46% |
| Cell-20 | 52% | 73% |
| Cell-21 | 42% | 56% |
| Cell-22 | 46% | 77% |
| Cell-23 | 53% | 72% |
| Cell-24 | 54% | 78% |
| Cell-25 | 32% | 96% |
| Cell-26 | 31% | 90% |
| Cell-27 | 40% | 75% |
| Cell-28 | 29% | 69% |
| Cell-29 | 28% | 60% |
| Cell-30 | 36% | 72% |
| Cell-31 | 37% | 71% |
| Cell-32 | 41% | 61% |
| Cell-33 | 48% | 75% |
| Cell-34 | 46% | 84% |
| Cell-35 | 46% | 68% |
| Cell-36 | 62% | 55% |
| Cell-37 | 49% | 88% |
| Mean | 42% | 72% |
| SD | 9% | 11% |

**Figure 6.**
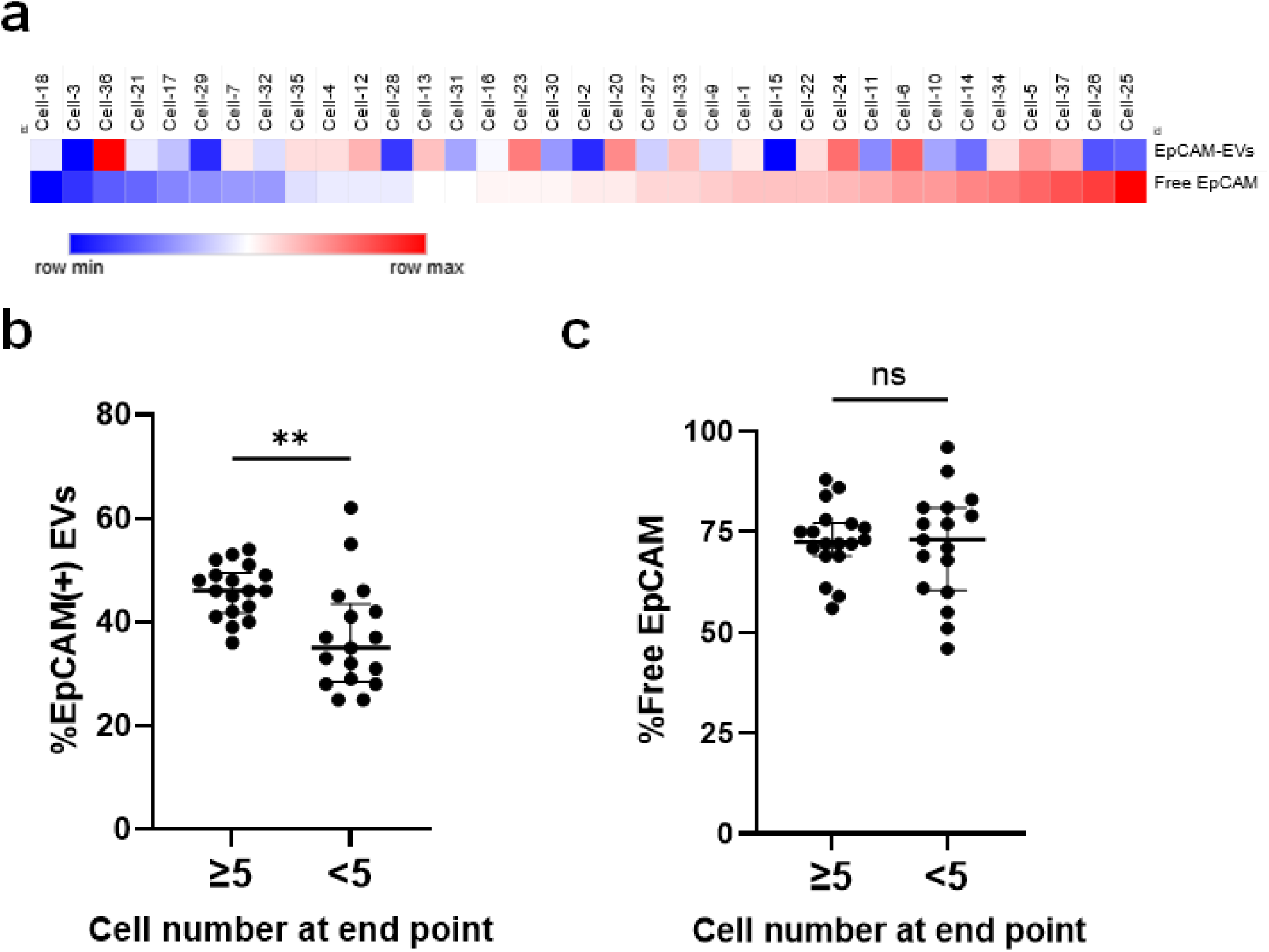
Quantification of EV marker-EpCAM co-expression and free EpCAM levels. **a. Heatmap of EV marker-EpCAM co-expression and free EpCAM levels**. Expression of EV markers (CD9, CD63, or CD81) and EpCAM in positive wells after thresholding was detected by high-resolution fluorescence microscopy, and EV marker–EpCAM co-expression levels and free EpCAM levels were quantified using CellProfiler™ 4. Percentages of EV marker–EpCAM co-expression (among total EVs) and free EpCAM (among total EpCAM signal) were transformed into z-scores, which were used for heatmap visualization in Morpheus (Broad Institute, https://software.broadinstitute.org/morpheus). **b. Comparison of EV marker–EpCAM co-expression percentages between wells containing ≥5 cells and wells containing <5 cells, based on endpoint cell counts after culture.** **c. Comparison of free EpCAM percentages between wells containing ≥5 cells and wells containing <5 cells, based on endpoint cell counts after culture**. Statistical significance was assessed using the Mann– Whitney test (P < 0.01; ns, P > 0.05).

## Discussion

We developed a microfluidic platform to analyze extracellular vesicles (EVs) released by cells derived from a single parental cell, enabling EV profiling from a specific, individual lineage. The cell array contains 17,305 wells designed to trap single cells and support subsequent culture after assembly with an EV array using a custom-fabricated housing. The EV array captures EV-associated signals originating from the trapped cells (i.e., clonal progeny from one parental cell). As a proof-of-concept application of the platform, we analyzed EV marker co-expression patterns and measured the level of EpCAM-positive EVs.

Two non-mutually exclusive conceptual models are often discussed to explain EV release and cargo composition.^1,5^ The “garbage bag” model posits that EV secretion can contribute to cellular homeostasis by exporting excess, obsolete, or otherwise undesirable constituents.^1,2,7–9^ In parallel, a “passive loading” model suggests that EV cargo composition may partly arise from probabilistic sampling of cellular components during vesicle biogenesis, such that abundant molecules are more likely to be packaged, thereby mirroring cellular signatures.^1,4^ In this study, our results appeared to be consistent with predominantly passive, abundance-driven cargo loading for the markers examined.

The abundance of EpCAM-positive EVs was positively correlated with the number of cells at the experimental endpoint. Because each well initially contains a single trapped cell, the endpoint cell number represents proliferative phenotype. Thus, this result suggests that more proliferative PC3 cells release higher levels of EpCAM-positive EVs. In contrast, the fraction of free EpCAM (i.e., non–EV-associated EpCAM) within the total EpCAM signal did not show a correlation. Together, these findings may indicate that proliferative cells release more EpCAM overall and that a substantial portion of this increased EpCAM is exported in an EV-associated form. This observation is consistent with previous findings that EpCAM expression can be elevated in proliferative cell states.^19–22^

EV marker co-expression profiles were heterogeneous. Because the tested cells were from the PC3 cell line, baseline cellular heterogeneity is expected to be relatively limited; nevertheless, substantial variability in EV marker co-expression was observed even among single-cell-derived clones within the same cell line. This may reflect state drift during clonal expansion over a relatively short culture period: given a PC3 doubling time of approximately 18 hours and a total post-trapping culture time of 72 hours, cells could undergo approximately 4 population doublings, during which molecular states may shift if EV cargo indeed reflects cellular status.

The developed microfluidic system enables EV analysis at near single-EV resolution from cells originating from a single parental cell, thus it can be broadly applied to diverse cell types. With a 100 μm well size, most mammalian cells can be efficiently trapped and analyzed. The target protein cargo (e.g., EpCAM) can be readily expanded by changing the primary antibodies coated onto the EV array. In addition, the target EV RNA cargo (miRNA or mRNA) could potentially be profiled by integrating single-EV in situ hybridization strategies, such as molecular beacon–based assays, on the captured vesicles.^23,24^ This platform has potential as an efficient tool for interrogating EV release from individual-cell lineages, thereby addressing the challenges posed by heterogeneity in both EV and cell populations.

## Acknowledgement

The authors thank Huy Vo, administrator of the Johns Hopkins University (JHU) WSE Whitaker Microfabrication Lab, for his consultation on the photolithography process. This work was supported by the Basic Science Research Program through the National Research Foundation of Korea, funded by the Ministry of Education [grant number 2021R1A6A3A14043146], the 2023 Postdoctoral Research Accelerator Award, funded by the Johns Hopkins School of Medicine to C.-J. Kim; the National Cancer Institute grant [grant number CA093900] to K. J. Pienta. The authors acknowledge the use of ChatGPT (OpenAI) for assistance with English language editing and improvement of grammar and readability. The authors reviewed and edited the output and take full responsibility for the content of this manuscript. The authors thank the current and past members of the Brady Urological Institute, especially members of the Pienta-Amend laboratory, for their critical feedback on this work.

---

**Supplementary Figure S1.**
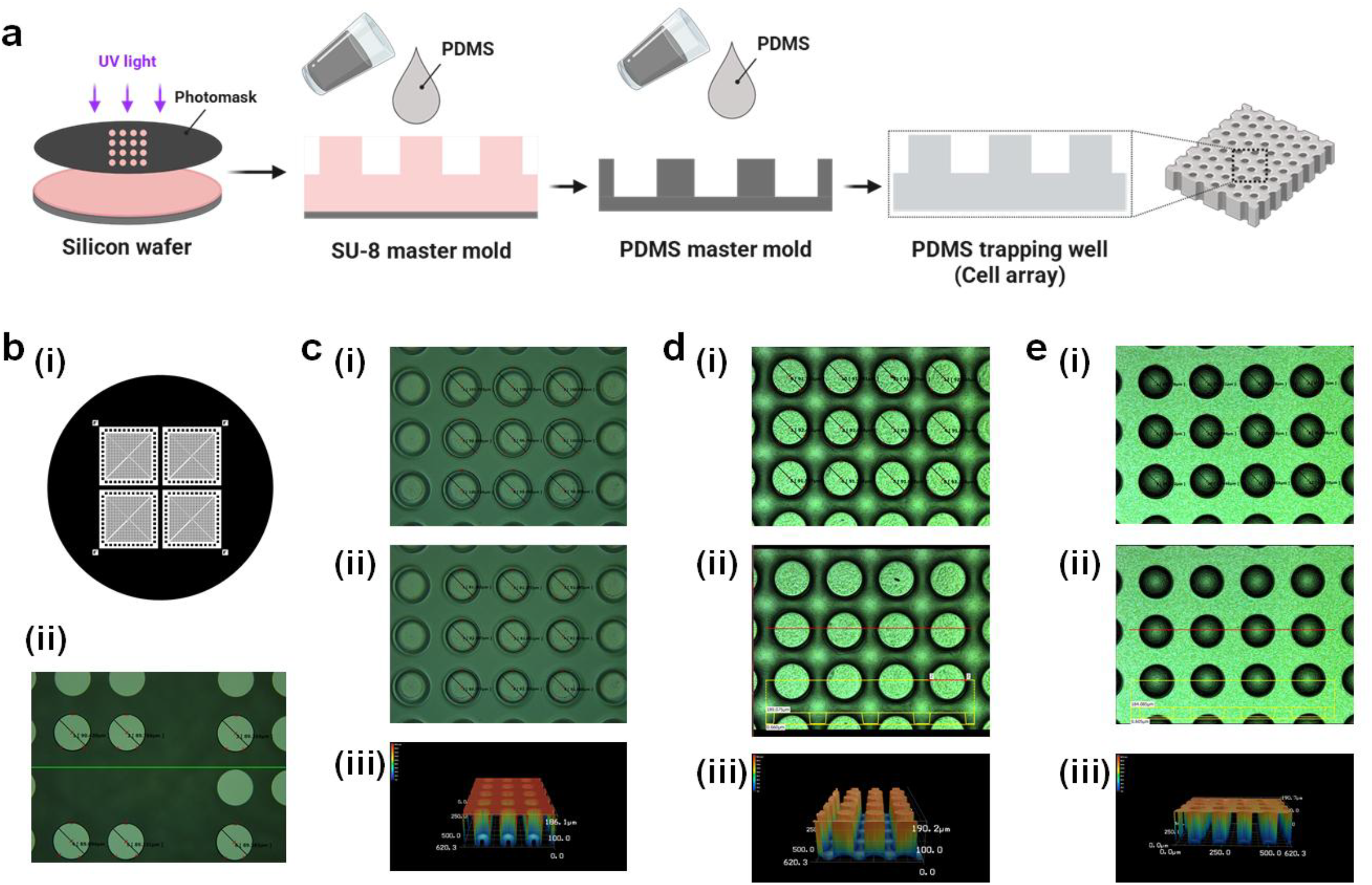
Fabrication of the cell-trapping well array (cell array). **a**. Brief schematic illustration of the fabrication procedure. An SU-8 microwell pattern was fabricated by photolithography to generate an SU-8 master on a silicon wafer. PDMS (base:curing agent = 10:1) was cast on the SU-8 master to produce a PDMS master mold (micropost pattern). PDMS (base:curing agent = 10:1) was then cast on the PDMS master mold to fabricate the final PDMS cell-trapping well array (cell array) using soft lithography. **b-e**. Stepwise validation of the fabrication procedure. **b**. (i) photomask design and (ii) digital microscope image of the photomask with diameter measurements. **c**. Characterization of the SU-8 master mold on a silicon wafer: (i) diameter measurement at the top surface, (ii) diameter measurement at the bottom surface, and (iii) 3D depth profiling. **d**. Characterization of the PDMS master mold: (i) digital microscope image with diameter measurements, (ii) depth profile with the corresponding top-view image, and (iii) 3D depth profiling. **e**. Characterization of the PDMS cell-trapping well array (cell array): (i) digital microscope image with diameter measurements, (ii) depth profile with the corresponding top-view image, and (iii) 3D depth profiling.

**Supplementary Figure S2.**
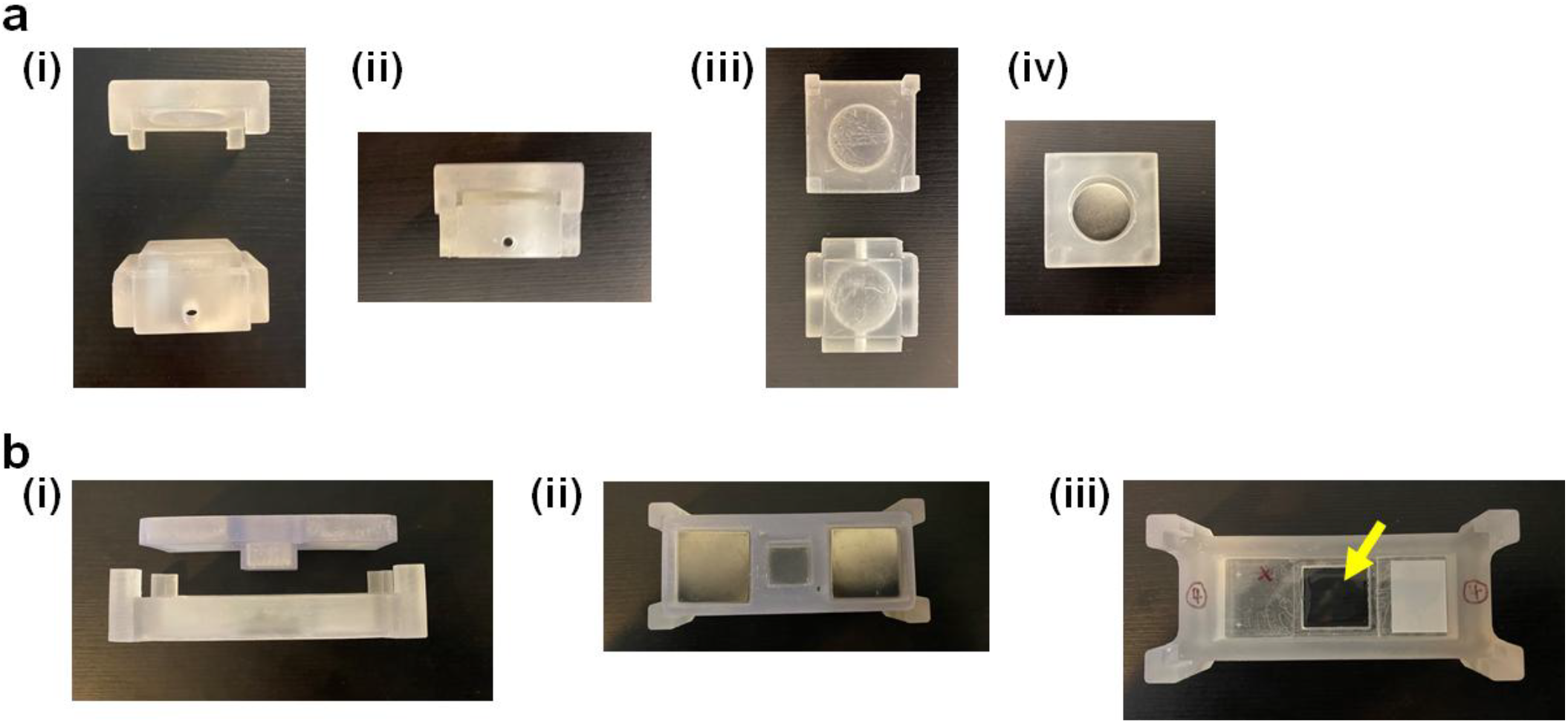
Photographs of the housing modules. **a. Compact housing**. (i) Cross-sectional view of the top and bottom parts shown separated. (ii) The housing in the assembled (housing-only) state. (iii) Top view of the inner surfaces of the top and bottom parts. (iv) Top view of the housing with magnets. **b. Specialized (windowed) housing for time-lapse live-cell imaging**. (i) Cross-sectional view of the top and bottom parts shown separated. (ii) Top view of the housing with magnets after assembly. (iii) Bottom part of the windowed housing. The central square region (black) is a transparent window (yellow arrow), enabling imaging of cells through the window during culture in the assembled device.

**Supplementary Figure S3.**
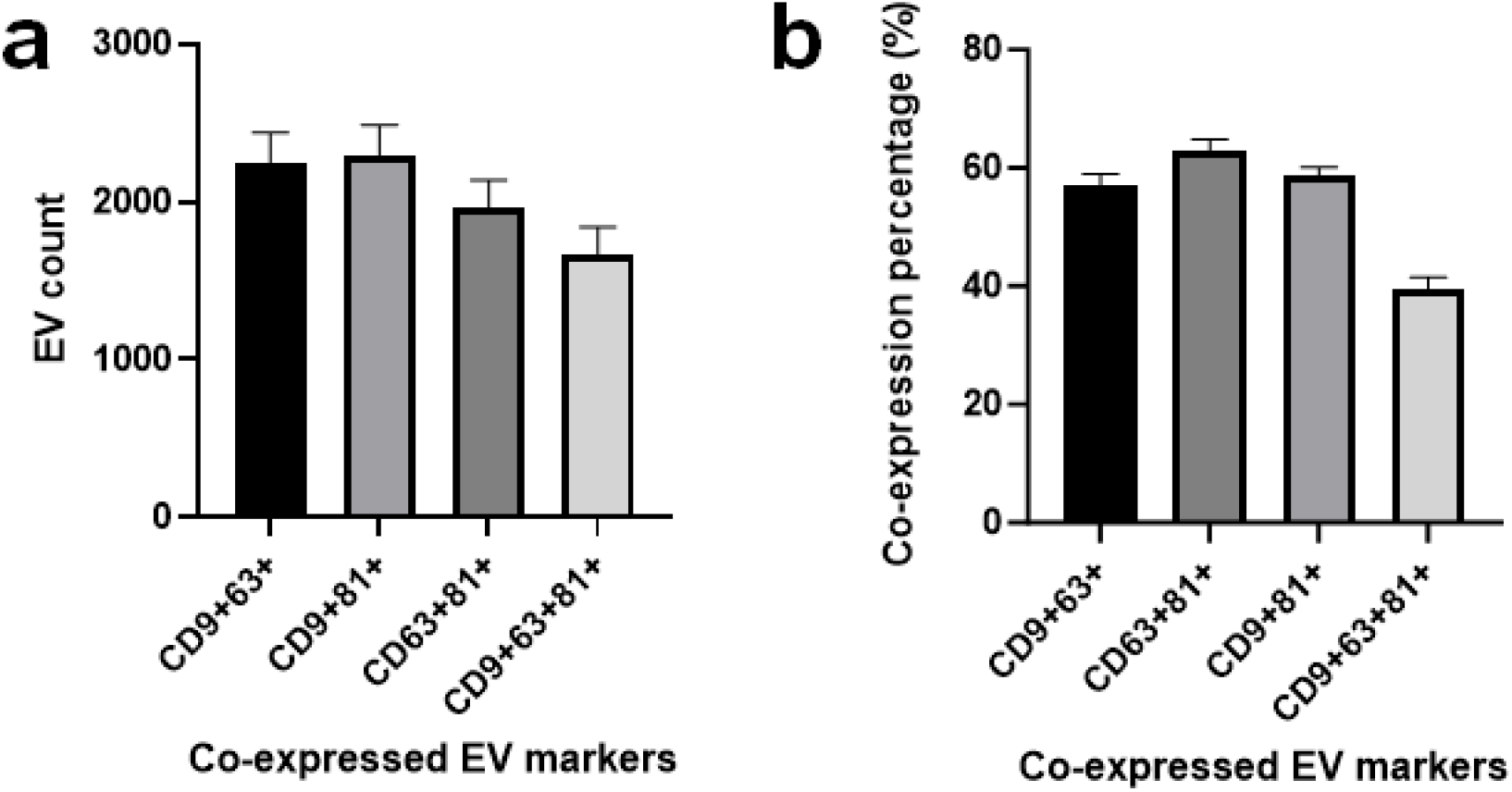
EV-marker co-expression analysis. **a**. Counts of EVs co-expressing EV markers. **b**. Percentages of EVs co-expressing EV markers.

**Supplementary Figure S4.**
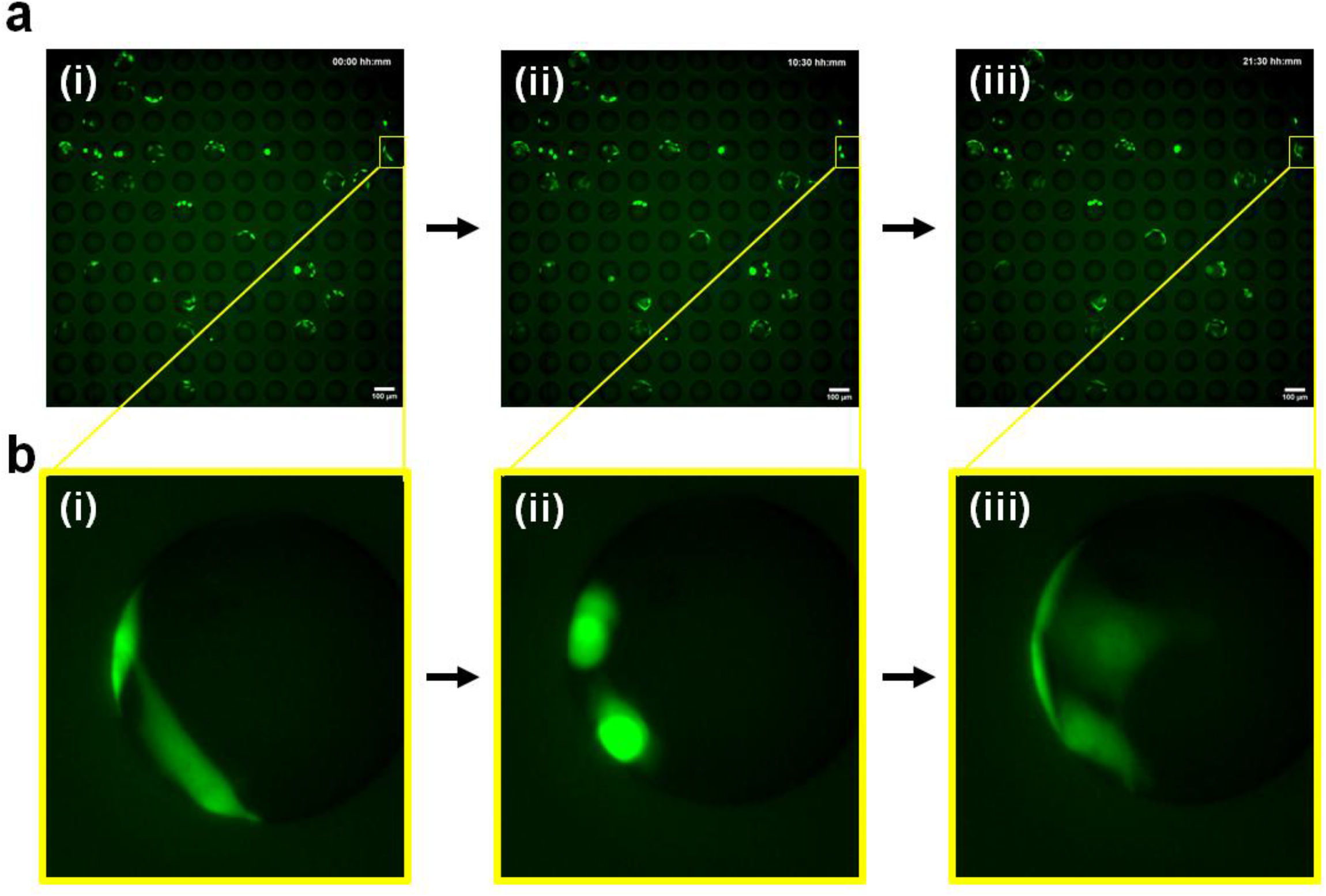
Snapshots from the cell array operation video. a. Snapshots of the cell-array operation video acquired at **(i)** 0 min, **(ii)** 10 h 30 min, and **(iii)** 21 h 30 min. b. A magnified view highlights a proliferating cell pair increasing from two to four cells over time: **(i)** 0 min, **(ii)** 10 h 30 min, and **(iii)** 21 h 30 min. PC3 cells were pre-labeled with 1 µM CellTracker™ CMFDA for 30 min and washed three times with complete medium prior to cell trapping.

**Supplementary Figure S5.**
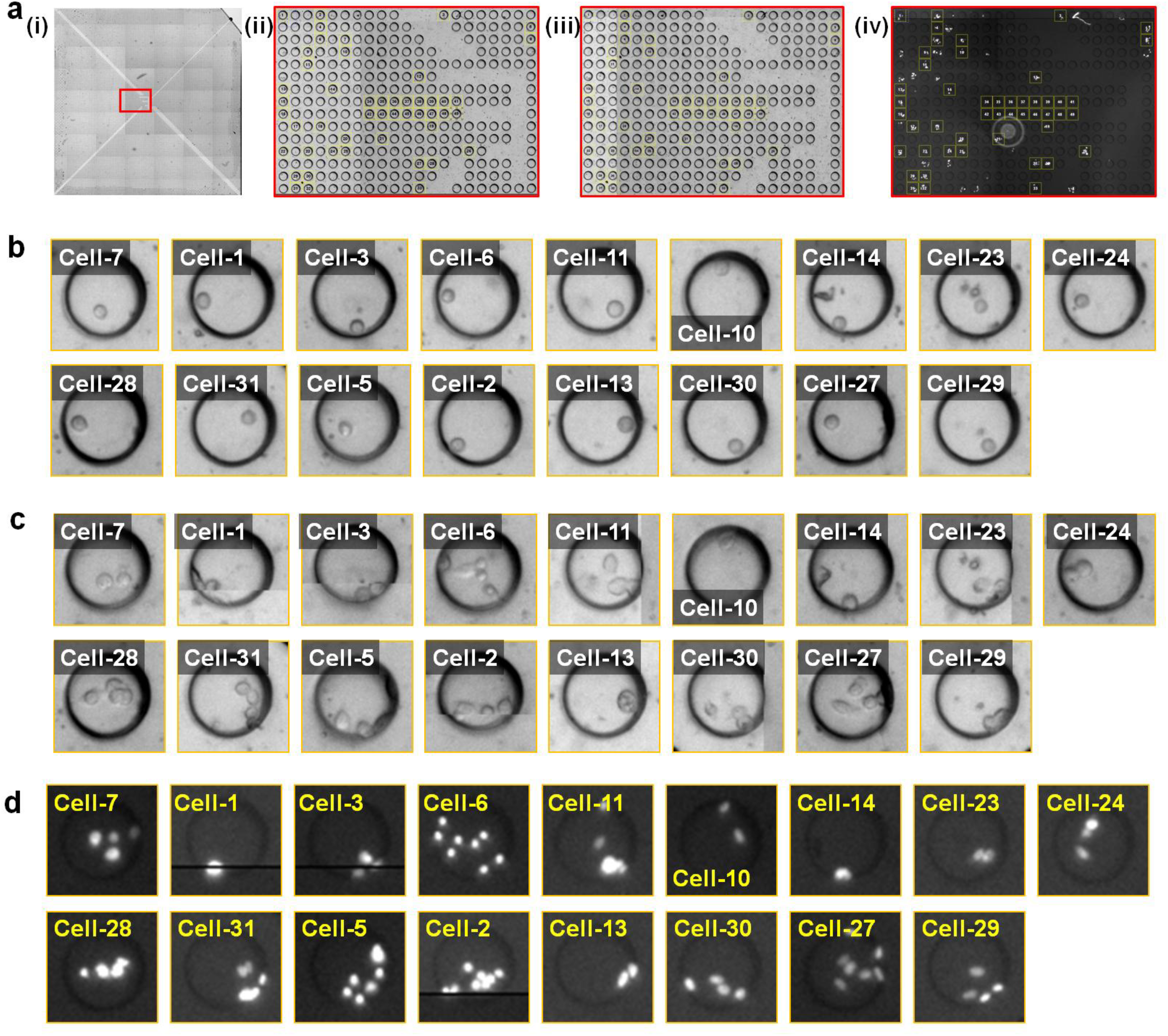
Cell-array wells analyzed for EV-marker expression. **a**. Images of the cell array and wells that passed the thresholding criteria for analysis. **(i)** Whole-field image of the cell array before housing assembly. Magnified views (red box) of **(ii)** Cell array after trapping. **(iii)** Cell array before housing assembly. **(iv)** Cell array after culture in the assembled device with nuclear staining (Hoechst). **b**. Images of wells containing a single trapped cell after trapping (cropped from panel **a-ii**). **c**. Images of the same wells before housing assembly (cropped from panel **a-iii**). **d**. Images of the same wells after culture (cropped from panel **a-iv**).

**Supplementary Figure S6.**
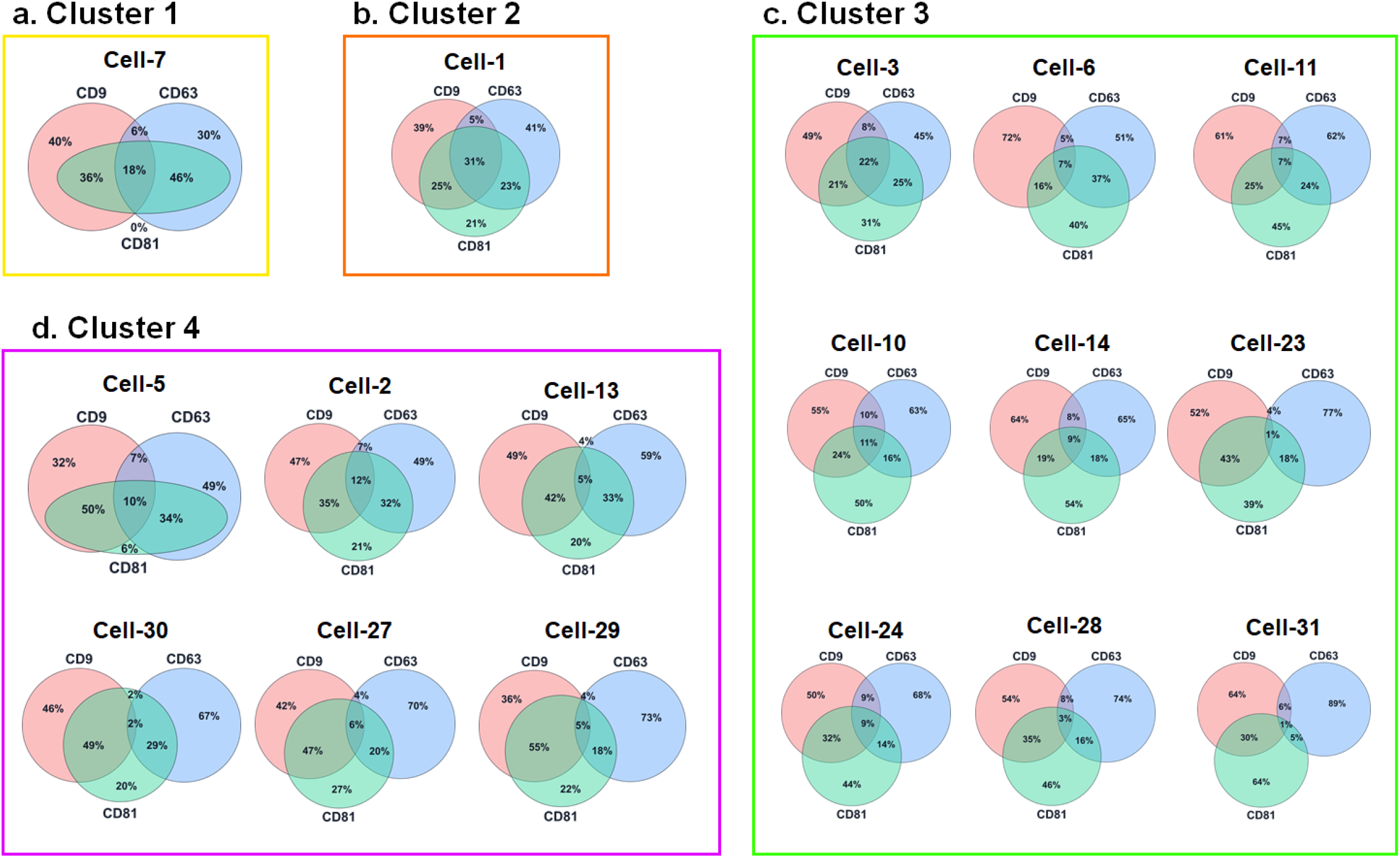
Venn diagrams summarizing EV-marker (CD9/CD63/CD81) co-expression for four clustering-defined groups.

**Supplementary Figure S7.**
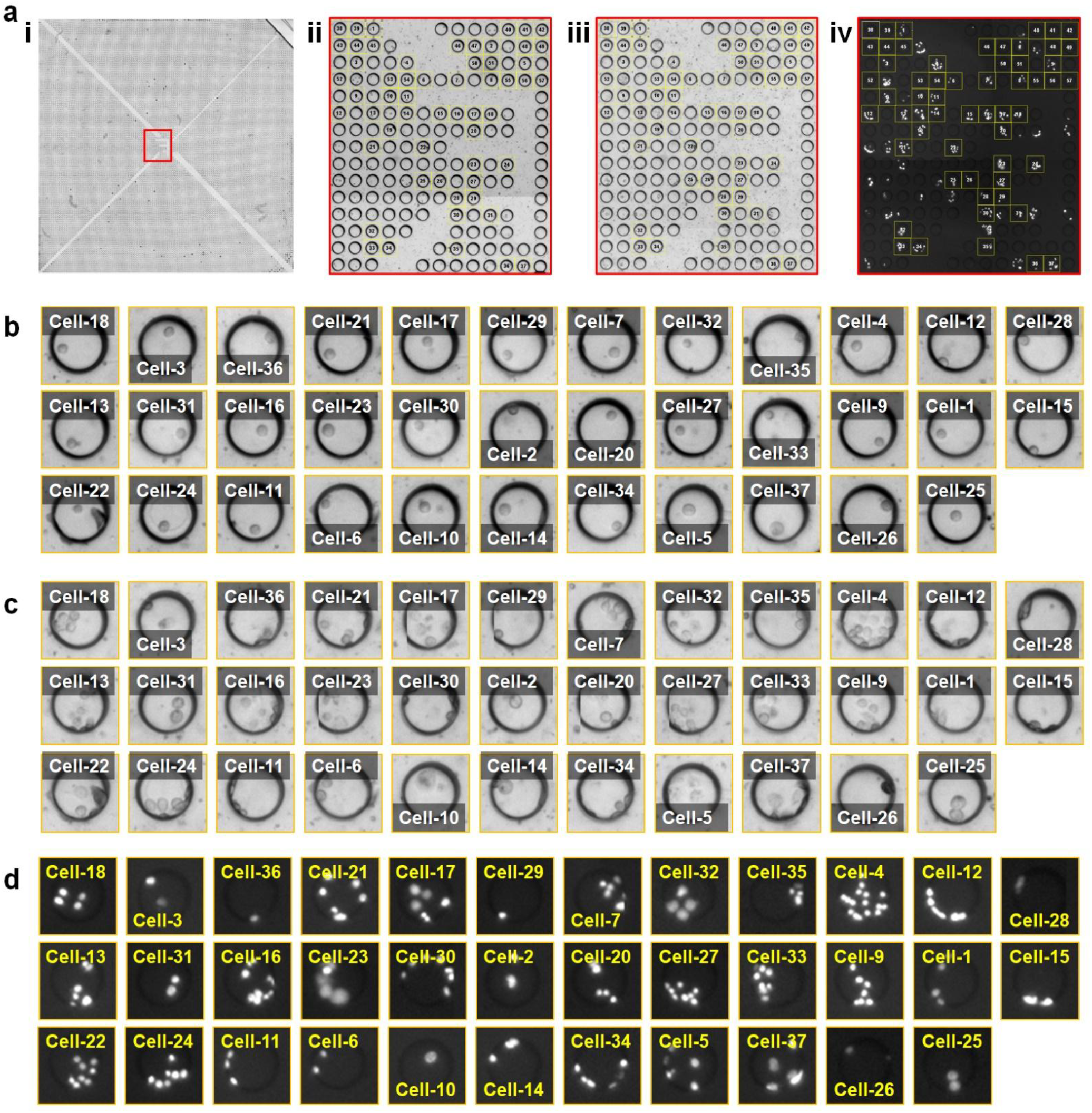
Cell-array wells analyzed for EpCAM-positive EVs and free EpCAM expression.. **a**. Images of the cell array and wells that passed the thresholding criteria for analysis. **(i)** Whole-field image of the cell array before housing assembly. Magnified views (red box) of **(ii)** Cell array after trapping. **(iii)** Cell array before housing assembly. **(iv)** Cell array after culture in the assembled device with nuclear staining (Hoechst). **b**. Images of wells containing a single trapped cell after trapping (cropped from panel **a-ii**). **c**. Images of the same wells before housing assembly (cropped from panel **a-iii**). **d**. Images of the same wells after culture (cropped from panel **a-iv**).

## Notes

### Competing Interest Statement

The authors have declared no competing interest.

